# Development of a Staining-based Electrophoretic Mobility Shift Assay for Analyzing Pbx1, DNA and HoxA9 Interactions

**DOI:** 10.64898/2026.06.03.729869

**Authors:** Kamal Rai, Ibukunoluwa Abigail Olaosebikan, Pegah Norouzi, Gabriella Pettipas, Abra G. Dadum, Madison Short, Kevin C. Courtney, Hacer Karatas Bristow

## Abstract

Transcription factors (TFs) are master regulators of gene expression and control a wide range of cellular functions including embryonic development, signaling pathways, immune response, and differentiation. A tight regulation of gene expression is crucial during all stages of life and changes in the TF function can lead to developmental abnormalities, diseases such as cancer, or resistance to treatment. Therefore, TFs are a promising class of drug targets, and the techniques that would contribute to the development of TF modulators are critical. Electrophoretic Mobility Shift Assay (EMSA) has been a primary tool to verify protein-DNA interactions, where a fluorescent, biotin, or isotope end-labelled DNA probe is used to quantify binding. Such labeling techniques, however, can be costly, time consuming, possess safety hazard risks and require capital equipment for imaging. Here, we optimized a label-free, staining-based EMSA to characterize a potential drug target, homeodomain (HD) TF Pre-B-cell leukemia homeobox-1 (Pbx1) and its binding partner Homeobox A9 (HoxA9). Staining the polyacrylamide EMSA gel with a DNA intercalating green cyanine dye – SYBR safe – allowed the visualization of Pbx1 homeodomain interactions with DNA at nanomolar concentrations and enabled quantitative determination of protein-DNA apparent binding affinity in the sub-micromolar range. Furthermore, a ternary complex of homeodomains of Pbx1 with HoxA9 and the DNA was also visible in the assay. We have shown that the staining-based EMSA can be used to evaluate inhibitors of Pbx1 that block the interaction with DNA. We have further validated the data from our assay with fluorophore labeling-based EMSA. Overall, using HD transcription factors Pbx1 and HoxA9 as a model, we have optimized a reliable and cost-effective staining-based EMSA that enables the high-sensitivity visualization and quantitative evaluation of transcription factor-DNA complexes without the need for end-labeled DNA probes. The streamlined workflow could be readily adapted to other DNA-binding proteins to study their interactions with the DNA, inhibitors, and other proteins.

## Introduction

TFs regulate gene expression by recognizing specific DNA binding elements and forming protein-protein interactions to recruit co-activators. Tight regulation of gene expression is crucial during all stages of life and changes in the TF function can lead to developmental abnormalities, diseases or resistance to treatment.^1^ While TFs are potential drug targets, they have historically been regarded as *undruggable* due to their large, disordered regions and lack of well-defined pockets.^2,3^ On the other hand, recent advances in understanding protein-protein interactions, screening techniques (e.g. fragment-based approaches), and novel targeting strategies (e.g. targeted -protein degradation, -translocation, - posttranslational modification)^4–7^ have renewed attention on TFs as viable drug targets, further emphasizing the importance of readily available techniques to evaluate TF modulators in binding assays.

Historically, the electrophoretic mobility shift assay (EMSA) has been the most frequently used binding assay for TFs that quantitatively visualizes their interactions with DNA. In this technique, a TF together with the cognate DNA, usually a synthetic oligonucleotide, is loaded into a non-denaturing polyacrylamide gel and then subjected to electrophoresis. The DNA migrates fastest when free, and much slower upon binding to a protein; multimeric complexes migrate even slower.^8,9^ A radioisotope, biotin or fluorophore end-label is often used to track the movement of the oligonucleotide. Since these methods require labeled DNA probes, a high cost is associated with testing multiple DNA sequences. Furthermore, there are tag-specific drawbacks; radioisotope labelling possesses safety risks, short half-life, and requires lengthy imaging and capital equipment. While the use of a fluorophore or biotin is safe, the inclusion of a tag can hinder the protein-DNA interactions.^8–11^ Therefore, an EMSA technique that eliminates the need for a tag to visualize protein-oligonucleotide interactions would be ideal to trace TF and DNA interactions. Such dye-based methods have been reported using Ethidium bromide (EthBr) and cyanine dyes such as TOTO, YOYO and SYBR Safe. In these techniques, the gel is treated with a DNA-intercalating dye upon electrophoresis to ensure that the DNA-protein complex is not disturbed. While promising, these techniques have been limited to initial verification of interactions and lacked further optimization and application.^12,13^

Here we optimized a staining-based EMSA to evaluate a potential drug target, transcription factor Pre-B-cell leukemia homeobox-1 (Pbx1) and its binding partner Homeobox A9 (HoxA9), both of which are homeodomain (HD) TFs that function during embryogenesis, regeneration, and hematopoiesis. Upregulation of Pbx1 and HoxA9 has been frequently observed in liquid and solid tumors. Pbx1 was first identified as an oncogenic transcription factor in B-precursor leukemia as part of the E2A-Pbx1 fusion protein.^14,15^ Significantly higher levels of Pbx1 mRNA have been observed in invasive breast, ductal and lobular carcinoma tissues compared to normal tissues.^16^ Furthermore, Pbx1 has been reported to play a role in chemoresistance in prostate and recurrent ovarian cancers.^17,18^ In addition, HoxA9 upregulation has been observed in more than 50 % of acute myeloid leukemia (AML) cases and is a predictor for poor prognosis.^19,20^

HoxA9-dependent leukemogenesis is greatly accelerated by the cofactors as Pbx.^21,22^ Overall, Pbx1 and HoxA9 are potential targets in cancer therapy, and inhibitors that block the interactions mediating the formation of Pbx1, DNA and HoxA9 complexes could lead to new therapies.

With a motivation to develop a binding assay to evaluate probes that can inhibit Pbx1 complexes, here we have optimized a staining-based EMSA to visualize binary and ternary complexes of Pbx1HD with DNA and HoxA9HD. We have shown that Pbx1HD interactions with an unlabeled DNA oligonucleotide can be detected in the nanomolar range, which enables the determination of an apparent binding affinity of Pbx1HD to its cognate DNA. In addition, we have shown that this assay can clearly detect the formation of the binary HoxA9HD-DNA complex as well as Pbx1HD, DNA and HoxA9HD ternary complexes using the optimized conditions. We have further confirmed the robustness of the data from the staining-based EMSA with fluorophore labeling-based EMSA as well as a Fluorescent Polarization (FP) assay. Next, we have demonstrated that staining-based EMSA can be efficiently used to evaluate inhibitors of Pbx1 complexes. Small molecule T417 and HoxA9-derived peptides inhibited binary and ternary complexes of Pbx1HD in a dose-dependent manner. Overall, using HD transcription factors Pbx1 and HoxA9 as a model, we have optimized a staining-based EMSA that enables the visualization and quantitative evaluation of transcription factor-DNA complexes without the need for labeled probes. The streamlined workflow could be readily adapted to other DNA-binding proteins to study to their interactions with the DNA, inhibitors, and other proteins.

## Results

### Purification of Pbx1HD and HoxA9HD

HDs of Pbx1 and HoxA9 are comprised of three helices where the helix 3 of each HD makes extensive interactions with the DNA major groove (Figure 1A). HDs also contact the DNA minor groove using basic lysine and arginine residues located at their N-terminals. In addition, Pbx1 has a short fourth helix that enhances the binding affinity to DNA.^23,24^ HoxA9 uses a tryptophan containing pentapeptide motif at the immediate N-terminus of the basic residues, which mediates binding with Pbx1.^25^ Accordingly, we designed HD expression constructs for each protein that contains the essential DNA binding elements (Figure 1B).

**Figure 1.**
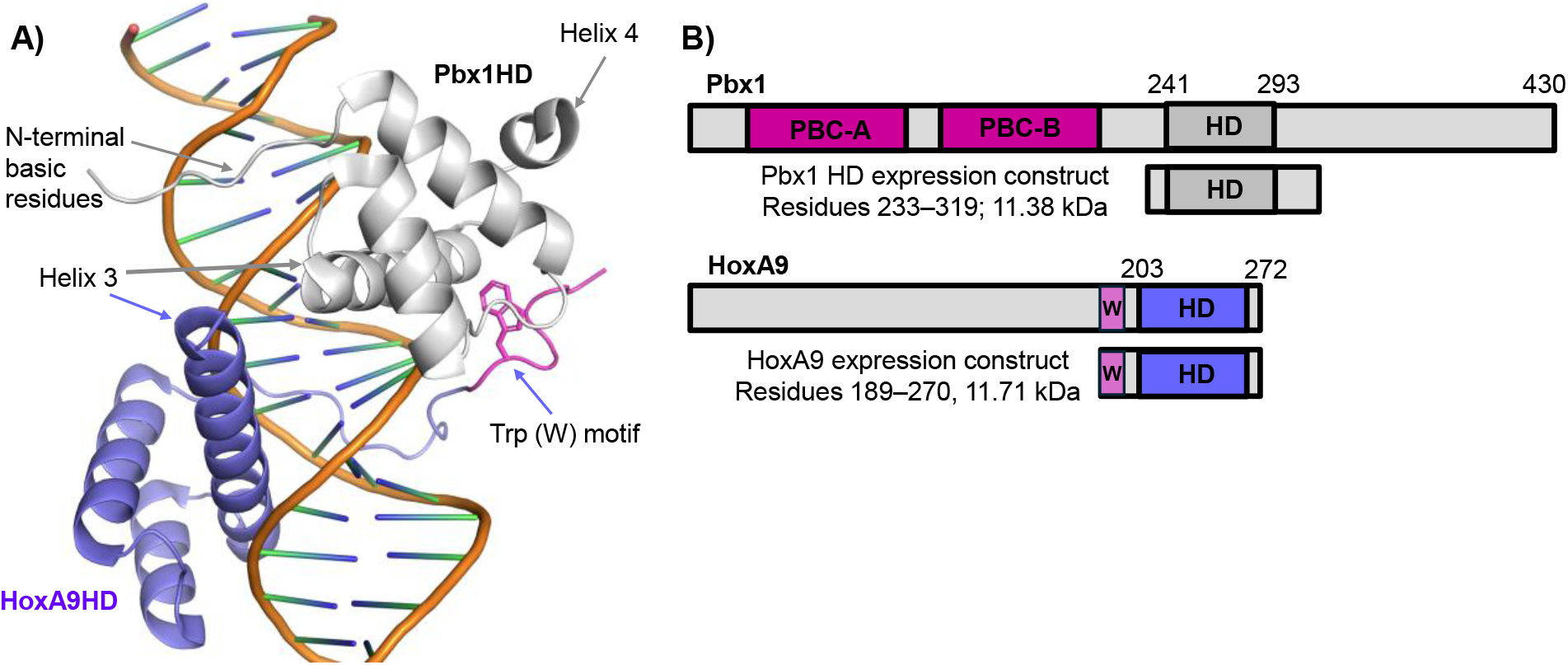
Structures of Pbx1 and HoxA9. A) Co-crystal structure of Pbx1HD, HoxA9HD and a DNA oligonucleotide (PDBID 1PUF). B) The domain structure of Pbx1, HoxA9, and the expression constructs used in this study.

Pbx1HD (233-319), comprised of N-terminal basic residues, helices 1-4 and a C-terminal His-tag, was expressed using lactose-mediated auto-induction conditions, followed by a two-step purification consisting of affinity (Figure 2A) and Size Exclusion Chromatography (SEC) (Figures 2B−2C). SEC analysis showed a single symmetric elution peak of about 14 mL, indicating a homogenous protein population (Figure 2B). A yield of 5 mg per 0.4L culture was obtained from expression in Terrific Broth (TB), which was nearly 5-fold more compared to expression in Luria-Bertani (LB) broth via IPTG induction. The Pbx1HD demonstrated high stability even during room temperature purification. Furthermore, it was soluble and stable not only with different buffer systems (e.g. MES pH 5.0 and 6.5, Tris pH 8.0 and 8.8), but also in deionized water as reported earlier.^26^

**Figure 2.**
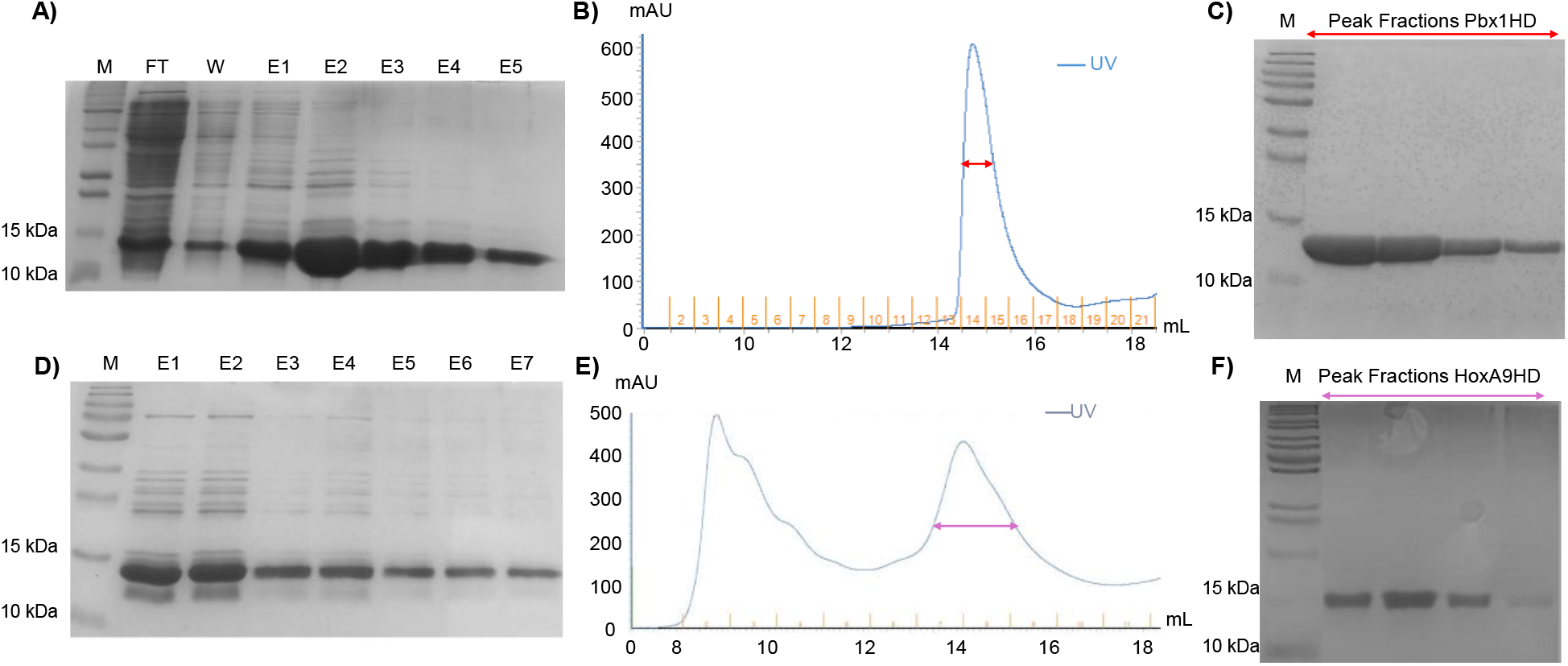
Two-step purification of A-C) Pbx1HD (11.38 kDa) and D-F) HoxA9HD (11.71 kDa). A and D: SDS Page images of affinity purification of Pbx1HD and HoxA9HD, respectively. B and E: Size exclusion chromatograms indicating the collected peak. C and F: SDS Page analysis of peaks collected during SEC. M: Marker, FT: Flow Through, W: Wash, E: repeated elution with 300 mM imidazole. Images were minimally processed for clarity and visibility. The original unedited images are provided in Figure S1.

HoxA9HD (189-270), which contains the pentapeptide motif, basic residues, helices 1-3 and a C-terminal His-tag, was expressed in LB medium via either IPTG or auto-induction, followed by two-step purification, as was used for Pbx1HD, yielding 1 and 2.9 mg of HoxA9HD, respectively, per 0.4L culture (Figure 2D−F). A broad retention peak around 15 mL via SEC (Figure 2E) was confirmed for size and purity (Fractions 7-10, Figure 2F).

In the label-free staining-based EMSA method optimized here, the DNA was loaded and run through an 8% polyacrylamide gel by electrophoresis. Then the gel was stained with DNA intercalating dye SYBR safe for 30 mins followed by washing briefly with water and imaging fluorescence (Ex/Em: epi-blue LED (460−490) nm / 518−548 nm. The wash step with water is essential as washing with a buffer gave significant noise and was therefore avoided (result not shown).

We initially assessed the sensitivity and the linearity of the signal at varying concentrations of a 20-base pair (bp) unlabeled DNA oligonucleotide that contained both Pbx1 (<u>TGAT</u>) and HoxA9 (<u>TTACG</u>) binding motifs (Table 1). In parallel, we tested a FAM (Fluorescein amidite)-labelled oligonucleotide with the same sequence as a control. We found that increasing concentrations of FAM-labeled DNA (5-500 nm) resulted in a linear increase in the signal (Figure 3A). The unlabeled DNA also yielded a concentration-dependent linear increase in the signal (Figure 3B). To assess the sensitivity, we lowered the unlabeled DNA concentration to 5-100 nM and increased the exposure duration during imaging (Figure 3C). The DNA bands remained detectable across the entire concentration range including 5 nM (0.6 ng) of the unlabeled DNA showing a high sensitivity.

**Table 1.**
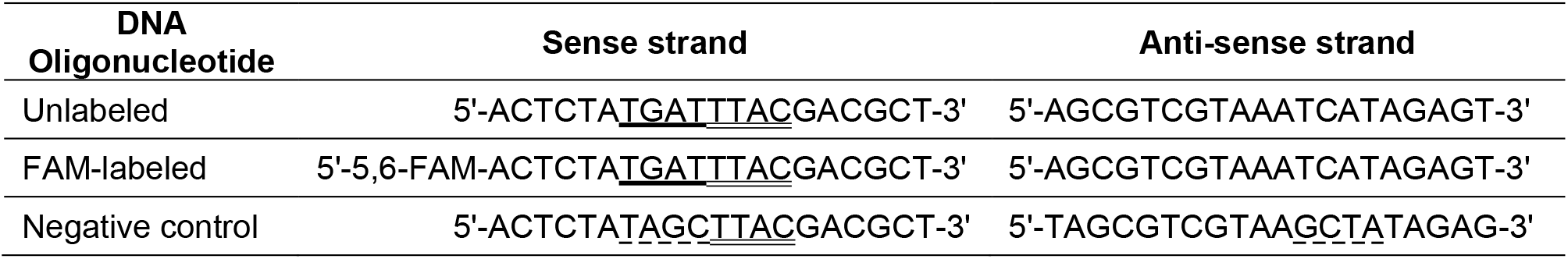
Sequences of the oligonucleotides used in the study. Cognate binding sequences for <u>Pbx1</u> and <u>HoxA9</u> are underlined. Pbx1 cognate sequence <u>TGAT</u> was mutated to <u>TAGC</u> in the negative control.

**Figure 3.**
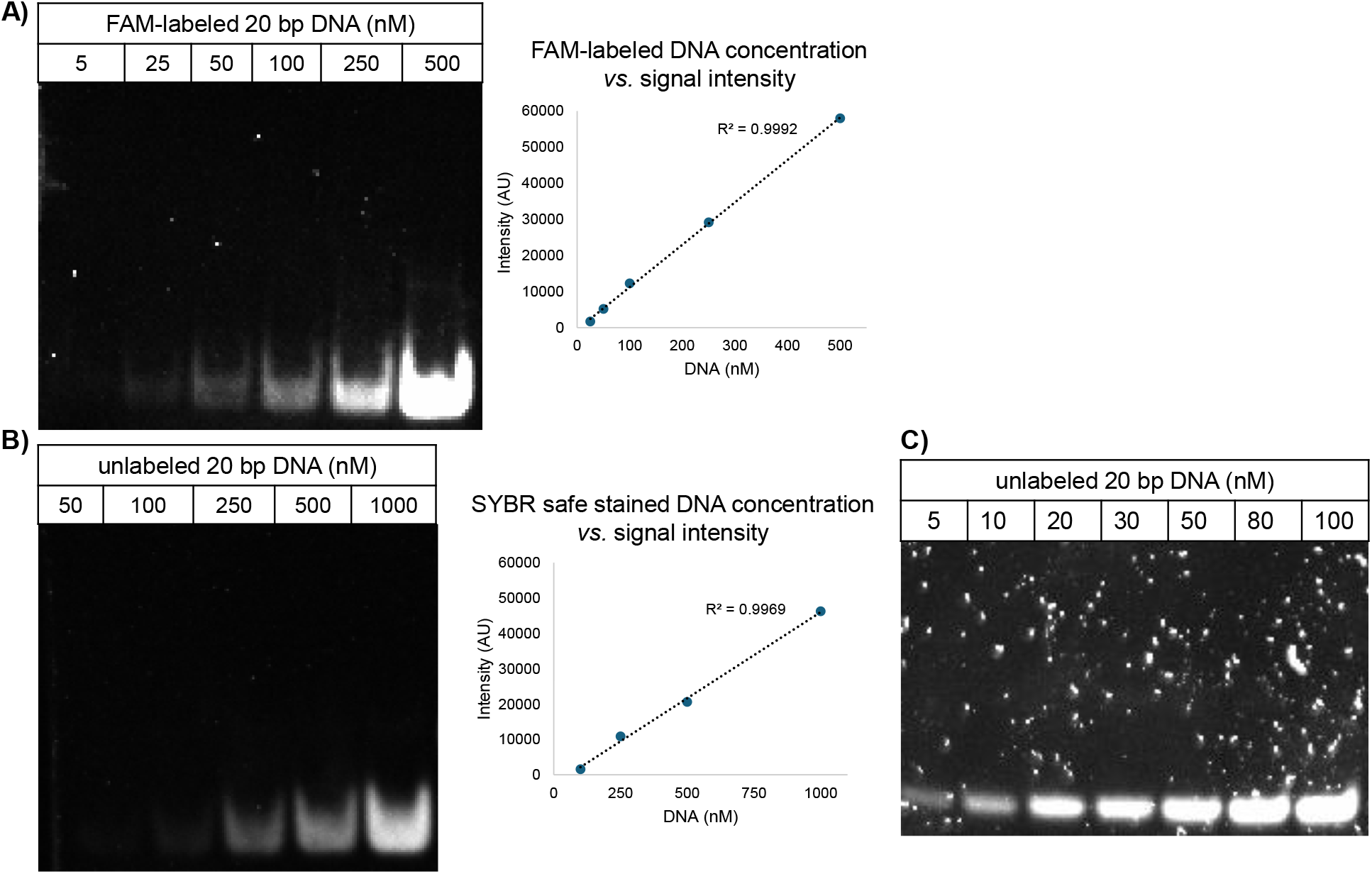
The detection sensitivity with 20 bp DNA oligonucleotides carrying the same sequence A) with a FAM-label, and B-C) without a label. The signal intensities in A and B were calculated using ImageJ and were plotted against the DNA concentration on the right.

Next, we tested if the Pbx1HD complex with the same unlabeled DNA oligonucleotide could be visualized with EMSA. When 25 nM of the DNA was titrated with increasing concentrations (0.1−1.8 µM) of Pbx1HD, we observed an increase in the signal intensity of the DNA shift, corresponding to the Pbx1HD-DNA complex, accompanied by a decrease in the free DNA signal intensity (Figure 4A). While 10 nM of the free DNA was visible, as shown in Figure 3C, the mobility shift with Pbx1HD was rather faint (result not shown). Therefore, 25 nM (3.06 ng) of DNA was used to visualize the Pbx1HD-DNA interactions. Another titration experiment with 5 nM of FAM-labelled DNA gave similar size DNA shift signals where the Pbx1HD-DNA complex was visible from 0.2 µM of Pbx1HD (Figure 4B). This indicates that FAM exhibits a 3-fold lower detection limit compared to staining-based method where the Pbx1HD-DNA complex was visible at 0.6 µM of Pbx1HD (Figure 4A).

**Figure 4.**
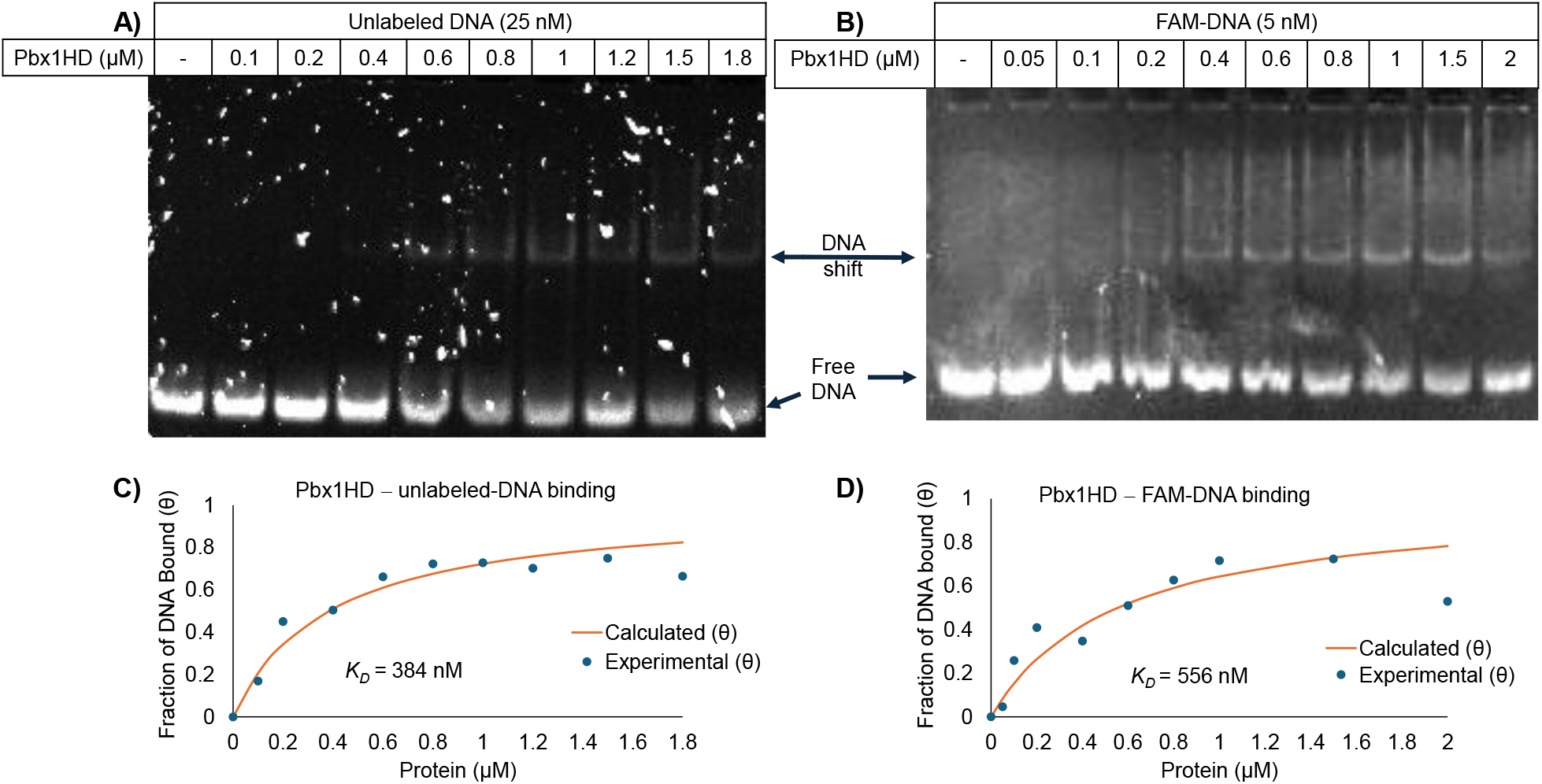
EMSA sensitivity to image Pbx1HD-DNA complex. The oligonucleotides titrated against Pbx1HD with A) unlabeled DNA, B) FAM-labelled DNA. Graphs showing the binding curve with their apparent *K*_*D*_ values of Pbx1HD to C) unlabeled DNA D) FAM-labeled DNA. The free DNA signal on each EMSA gel, as calculated using ImageJ, subtracted from no protein control to determine the fraction of DNA bound (θ) and was plotted against the protein concentration. The apparent *K*_D_ values were determined using Langmuir’s isotherm.

The imaged gels in Figures 4A and 4B were next used to calculate the apparent binding affinity (*K*_D_) of Pbx1HD to DNA. The free DNA signal on each well, as calculated using ImageJ, was subtracted from the no protein control to determine the fraction of DNA bound (θ) and was plotted against the protein concentration and the resulting binding curves were fitted to Langmuir binding isotherm (Figures 4C and 4D). Both EMSA techniques, i.e. using unlabeled and FAM-labeled DNA, gave similar *K*_D_ values which were 384 nM and 556 nM, respectively. The biological replicates also yielded consistent *K*_D_ values (Figure S2 and S3). To further evaluate these results, the FAM labelled DNA was tested using a fluorescence polarization (FP) assay which gave a *K*_D_ of 84 ± 19 nM (Figure S4).

Based on the initial Pbx1HD-DNA binding experiments (Figure 4A), we have chosen 0.5 µM of the Pbx1HD for future experiments as the DNA shift (i.e. Pbx1HD-DNA complex) signal is clear and thus inhibition of the complex would be easily visualized. Furthermore, the DNA shift signal becomes more smeared - an indication of oligomeric heterogeneous complexes - with concentrations above 1.5 µM of Pbx1HD, and therefore such high protein concentrations are not suitable for EMSA in our experimental setup.

To show that the DNA shift with the Pbx1HD is due to a specific interaction, a negative control, where the cognate DNA sequence of Pbx1HD <u>TGAT</u> mutated to <u>TAGC</u> (Table 1), was tested. No visible DNA shift band in the mutant DNA lane with Pbx1HD was observed (Figure 5A). To further evaluate the specificity, we addressed whether the His-tag on Pbx1 interferes with the observed DNA interaction. Accordingly, a 6-His peptide was tested in the staining-based EMSA and the FP assay. No competitive inhibition was observed in either the EMSA (Figure 5B) or the FP-assay (Figures 5C), validating that the Pbx1HD binding to its cognate DNA sequence is specific.

**Figure 5.**
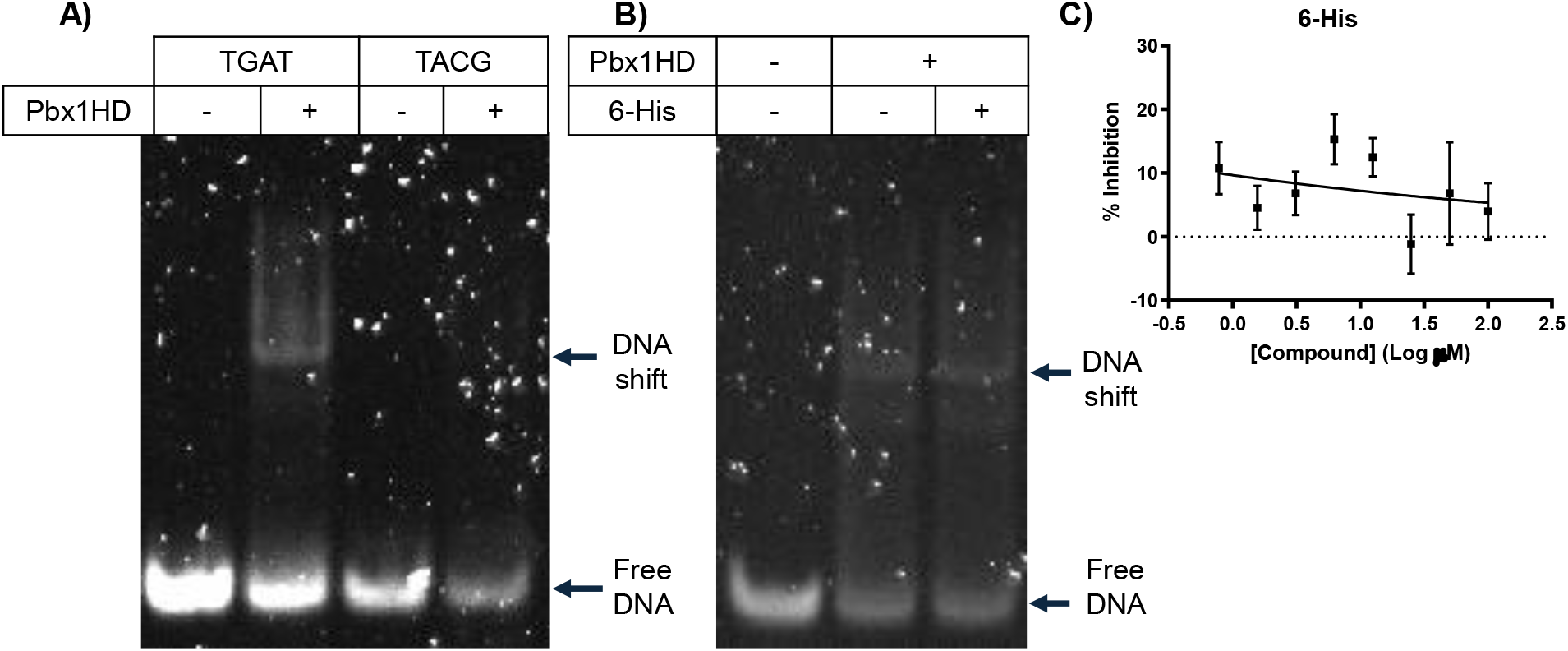
EMSA selectivity to image Pbx1HD–DNA complex. A) Binding of Pbx1HD to the DNA oligonucleotide with cognate TGAT sequence *vs*. a negative control oligonucleotide where TGAT is mutated to TACG. Hexa-Histidine peptide (6-His) did not interfere Pbx1HD and DNA binding as shown with B) EMSA, and C) FP-based competitive binding assay. A concentration of 0.5 µM of Pbx1HD, 25 nM (TGAT) and 15 nM (TACG) oligonucleotides were used for the EMSA experiments.

We next applied the staining-based EMSA to visualize the HoxA9HD complex with the same 20 bp oligonucleotide used for Pbx1HD. This oligonucleotide contains the cognate HoxA9 binding sequence <u>TTACG</u> (Table 1) and is expected to form a complex with HoxA9HD. Consistently, increasing DNA shift signals corresponding to HoxA9HD-DNA binary complex and diminishing free DNA signals were obtained as the protein concentration increased (Figure 6A).

**Figure 6.**
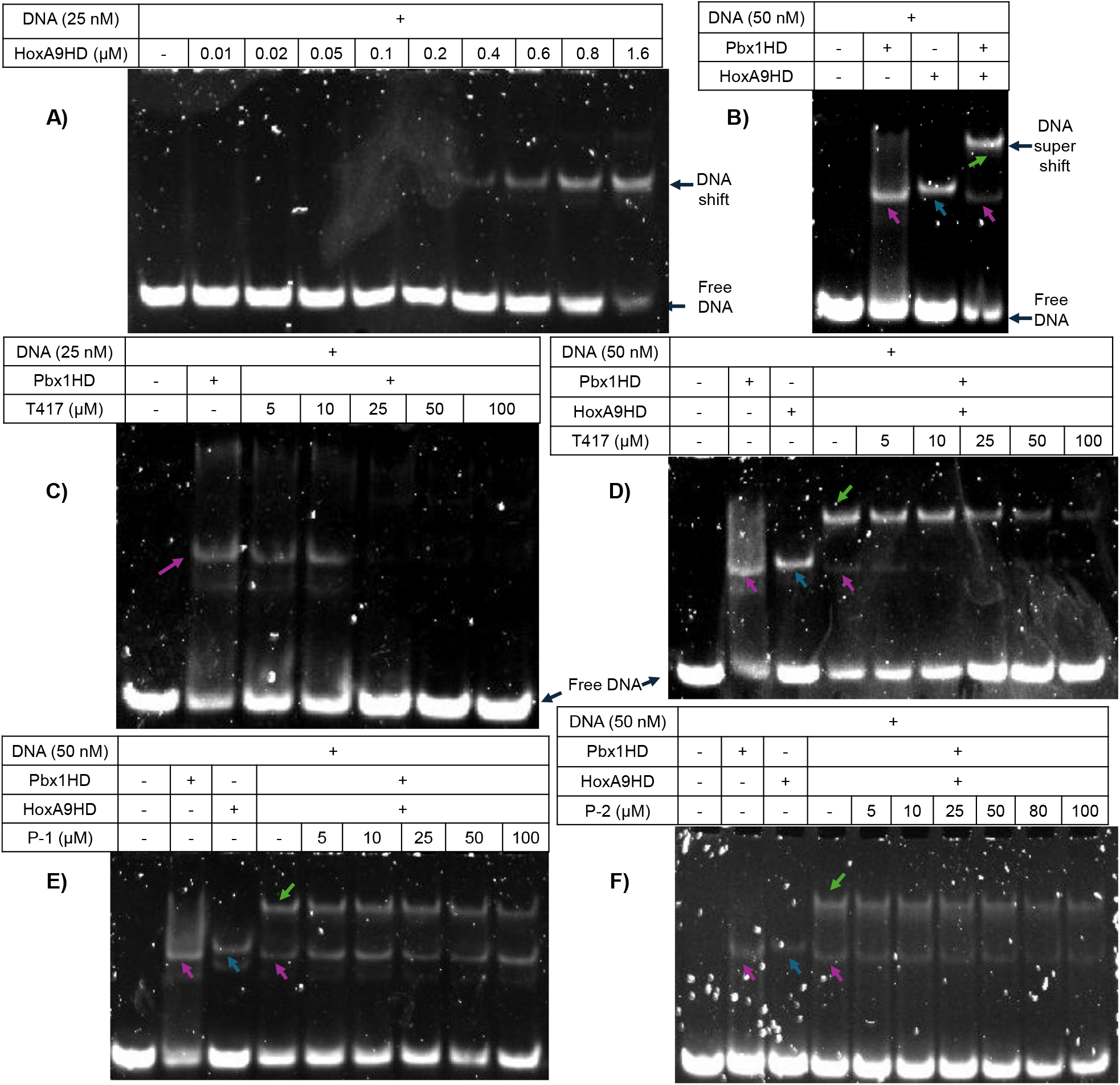
HoxA9HD and Pbx1HD complexes with DNA as visualized with the staining-based EMSA. Colored arrows represent the binary complexes of Pbx1HD-DNA (pink), HoxA9HD-DNA (blue) and ternary complex of Pbx1HD-DNA-HoxA9HD (green). A) Formation of HoxA9HD complex with the unlabeled DNA. B) Ternary DNA complex of Pbx1HD (0.5 µM) and HoxA9HD (0.5 µM) as shown with green arrow. Inhibition of C) Pbx1HD-DNA complex, and D) ternary complex with small molecule inhibitor T417. Depletion of the complex signal and increase in the free-DNA signal are indicators of inhibition. E-F) Inhibition of the ternary complex with peptide inhibitors P-1 and P-2. A concentration of 0.5 µM of each protein was used for all the inhibition (C-F) experiments.

Encouraged by the clear visualization of binary DNA complexes of Pbx1HD and HoxA9HD in the staining-based EMSA, we assessed whether a ternary complex of Pbx1HD, HoxA9HD and the DNA would be visible. A clear super shift (green arrow) above the DNA shifts of binary complexes (pink and blue arrows) when HoxA9HD and Pbx1HD were mixed with the DNA showed that the ternary complex could be visualized with the staining-based EMSA (Figure 6B). A concentration of 50 nM DNA was chosen for the ternary complex experiments, because when 25 nM of the DNA oligonucleotide was used, while the ternary complex was visible (Figure S5), the band was rather faint and might not be as suitable to monitor the inhibition.

### Inhibition of Pbx1 complexes as determined with the staining-based EMSA

Upon imaging binary and ternary complexes of Pbx1HD, HoxA9HD and the DNA, we assessed the applicability of the optimized staining-based EMSA to test the effect of Pbx1 inhibitors. When Pbx1HD (0.5 µM) was titrated with 5 μM – 100 μM of T417, a small molecule Pbx1-DNA interaction inhibitor (SPR IC_50_ = 3.35 µM),^27^ we observed a concentration dependent depletion in the DNA shift of the Pbx1HD-DNA complex (pink arrow) accompanied with increasing amount of free DNA signal (Figure 6C). We next tested the same molecule, T417, to visualize inhibition of the ternary complex. We hypothesized that the molecules that block the interaction of Pbx1HD with the DNA would consequently inhibit the ternary Pbx1HD-DNA-HoxA9 complex. Consistently, T417 treatment reduced the ternary complex formation in a dose-dependent manner (Figure 6D), validating that the staining-based EMSA can accurately measure both binary and ternary complex assembly/disassembly.

Lastly, we tested a peptide derived from HoxA9 to block the ternary complex formation (P-1, Figure 6E). A moderate decrease in the intensity of the ternary complex band and an increase in the free DNA band indicated weakening of the ternary complex. These results suggest that the HoxA9 peptide P-1 can weakly suppress the ternary complex assembly. When testing a second peptide (P-2), inhibition of the ternary complex was accompanied by a decrease in the free-DNA, suggesting that P-2 binds to the DNA rather than Pbx1 to inhibit the ternary complex.

## Discussion

While TFs are a challenging class of therapeutic targets due to lack of well-defined binding pockets, recent advancements in novel targeting strategies have brought attention to TFs as novel targeted therapies.^2,4–6^ Therefore, accessible binding assays, such as EMSA, to evaluate TF interactions with modulators are critical to accelerate discovery efforts. Here, we established a cost-effective and accessible EMSA binding assay for transcription factors Pbx1 and HoxA9, both of which are potential drug targets.

We initially designed expression constructs for homeodomains of Pbx1 and HoxA9 that are critical for DNA binding. Both constructs contained N-terminal basic residues and helices 1-3 of the homeodomains that interact with the DNA minor grove and major groove, respectively. An additional C-terminal extension, a fourth helix, was also included in Pbx1HD as it enhances DNA binding. Affinity purification with cobalt resin yielded high purity preparations of both Pbx1HD and HoxA9HD. Pbx1HD expression in TB medium via auto-induction gave 5-times higher yield compared to the expression in LB medium via IPTG induction. Higher cell density in TB medium and the non-toxicity of the lactose could be attributed to the higher yield from TB with auto-induction.^28,29^ Pbx1HD demonstrated excellent stability at a range of pH and in water.^26^ Overall, we have shown the recombinant protein expression and purification of Pbx1HD and HoxA9HD with good-to-moderate yields, i.e. 5 mg/0.4L culture and 2.9mg/0.4L culture, respectively, which facilitated the subsequent staining-based EMSA optimization (Figure 2).

Conventional EMSA methods rely on radioisotope, fluorophore, or biotin-labeled oligonucleotide probes to visualize nucleic acid interactions with proteins. Although these approaches have been widely used, they present several drawbacks, including a high cost associated with testing multiple probes. Radioactive labeling requires specialized facilities and safety protocols, while fluorophore or biotin labeling may influence DNA-protein binding if the label is near the binding site.^10,11,30,31^ These limitations can significantly hinder successful evaluation of multiple DNA probes as well as potential inhibitors.

Staining-based EMSA strategies, which use unlabeled oligonucleotides and are stained after electrophoresis, have been reported, however these prior studies lacked optimization for quantitative estimation and relied on micromolar DNA concentrations for protein-DNA interactions.^12,32^ As such, these prior methods were not suitable for high-sensitivity experiments, including the characterization of TF-DNA inhibitors. In the current study, using HD transcription factors Pbx1 and HoxA9 as a model, we optimized a staining-based EMSA that enables the visualization and quantitative evaluation of transcription factor-DNA complexes without the need for labeled probes.

In the optimized technique described here, a native polyacrylamide gel is run with an unlabeled DNA oligonucleotide, then stained with DNA-intercalating dye SYBRSafe, followed by a brief wash step with water. We observed a linear correlation between the unlabeled DNA concentration and the signal. While a higher sensitivity appeared with the FAM-labeled DNA as seen in Figures 3A and 3B, the detection sensitivity with the unlabeled-DNA improved when sub-100 nM DNA concentrations were used and was similar to that was reported.^12^ While not reported here, it is plausible that the sensitivity would be further enhanced with longer oligonucleotides, as there will be a larger surface area for the DNA-intercalating dye that would yield much higher signal.

With the DNA detection sensitivity at nanomolar concentrations, the Pbx1HD-DNA apparent binding affinity determination was feasible. Quantifiable DNA shift signals upon titration with Pbx1HD were observed yielding an apparent *K*_D_ = 359 nM, which was similar to that of FAM-labeled DNA (*K*_D_ = 556 nM) in EMSA showing that staining-based EMSA can be as quantitative as fluorescent EMSA (Figure 4). The same FAM-labeled DNA, however, showed a 3-fold higher affinity (*K*_D_ = 84 nM) in the FP-assay (Figure S4). Such discrepancy is common, and could be due to the low stability of the complex in gel, fast k_off_ rates or the gel composition.^9,33^

In addition to detecting binary DNA complexes of Pbx1 and HoxA9, the optimized conditions enabled the visualization of higher order complexes with nanomolar concentrations of the DNA. It has been shown that Pbx1 and HoxA9 form a cooperative complex with DNA. HoxA9 binding to DNA is weak, and addition of Pbx1 enhances interaction with the DNA.^25,34^ Consistent with the literature, we observed the cooperative ternary complex (Figure 6B). The ability of the staining-based EMSA to resolve these complexes demonstrates its applicability for studying transcription factor cooperation and multiprotein assemblies on DNA.

The major objective of this study was to establish an EMSA assay that is convenient for evaluating modulators of Pbx1 complexes. Based on the initial binding experiments, we have chosen 25 nM and 50 nM of the DNA oligonucleotide for the binary and ternary complex inhibition experiments, respectively. Since there was a hook effect with both proteins (i.e., 1.5 μM for HoxA9 and at 1.8 μM Pbx1HD), a lower concentration was used. Ideally, the protein concentration for competition experiments should be chosen within the linear range in the binding curve. Accordingly, we used 0.5 µM of each protein for the competitive binding experiments.

Using the optimized conditions, we tested inhibitors targeting the Pbx1 interaction network. The small molecule inhibitor T417, shown to block the Pbx1HD interaction with a biotin labeled DNA probe,^27^ disrupted the Pbx1HD and unlabeled DNA binary complex in a dose-dependent manner, showing that the staining-based EMSA can be a reliable alternative to assess protein-DNA interaction modulators, and can therefore replace the labeled probe EMSA.

Considering the biological significance of Pbx1 and HoxA9 interactions to contribute to oncogenic transcriptional programs,^34^ we assessed if Pbx1 modulators, such as T417 and HoxA9 derived peptides, can interfere with the Pbx1HD-DNA-HoxA9HD ternary complex formation. The small molecule T417 suppressed the ternary nexus in a dose-dependent manner. In their original study, Shen et al. showed that T417 inhibits Pbx1 ternary complex formation with Meis1 and the DNA via blocking the Pbx1 interaction with the DNA.^27^ Here, we have shown an additional mechanism with T417; blocking Pbx1 interaction with DNA also disrupts complex formation with HoxA9 suggesting that inhibiting Pbx1-DNA interaction could be a viable strategy to target HoxA9.

Next, a HoxA9 derived peptide (P-1, structure not disclosed here) was tested to disrupt the ternary complex. Although not as effective as T417, this peptide also suppressed the ternary complex formation (Figure 6E). Another peptide, P-2 (structure not disclosed here), also inhibited the ternary complex along with a decrease in free DNA signals suggesting that P-2 might bind to the DNA rather than Pbx1 (Figure 6F). These findings demonstrate that the staining-based EMSA is a practical approach to develop and mechanistically evaluate transcription factor inhibitors.

While the method presented here offers several noted advantages, certain limitations should also be considered. As with conventional EMSA approaches, interpretation of binding affinities relies on equilibrium assumptions and gel conditions that may not fully reflect other higher sensitive approaches such as SPR, ITC or FP. In addition, DNA intercalating dyes could potentially influence the stability of protein-DNA complexes during staining, although the post-electrophoretic staining approach used in this study minimizes this risk.

In summary, using HD transcription factors Pbx1 and HoxA9 as a model, we have optimized a simplified, cost-effective, and reliable staining-based EMSA method that enables high-sensitivity visualization and quantitative analysis of transcription factor-DNA complexes without the need for labeled DNA probes. The assay successfully enabled detection of binary and ternary DNA complexes of Pbx1HD and HoxA9HD and can be used to mechanistically evaluate inhibitors targeting these interactions. Our approach provides a practical tool for studying transcription factor DNA complexes and may facilitate the discovery and characterization of modulators targeting transcription factor regulatory networks. The streamlined workflow could be readily adapted to other DNA-binding proteins to study their interactions with the DNA, inhibitors, and other proteins.

## Materials and Methods

Oligonucleotides (Integrated DNA Technologies), 6-His peptide (APExBio), TCRS-417 (Ambeed Inc) were commercially obtained. The oligonucleotides were resuspended in Tris (10 mM, pH 7.5), NaCl (150 mM), EDTA (1 mM) DTT (0.1 mM) to a concentration of 200 μM, aliquoted and stored at -20 °C. A 10 mM stock solution of the peptides and TCRS-417 were prepared in DMSO and stored at -20 °C.

### Plasmids and transformation

Lyophilized plasmids with cloned human Pbx1HD (233-319) and HoxA9HD (189-272) in pET 28b^+^ expression vector under the control of T7 promoter were obtained from GenScript. Each plasmid was resuspended in ddH_2_O to 40 ng/μl and used for transformation into chemically competent expression hosts *E. coli* (rosetta gami 2) and BL21 DE3 via heat shock method. Chemically competent *E. coli* DH5alpha cells were used for plasmid amplification.

### Recombinant protein expression

<u>Expression via autoinduction:</u> Cells were grown in TB or LB medium supplemented with α-lactose monohydrate (10 mg/mL), MgSO_4_ (2 mM) and kanamycin (50 μg/mL) for 4 hrs at 37 °C. Then, the temperature was reduced to 25 °C and the culture was incubated for another 22h before harvesting. <u>Expression via IPTG induction:</u> Cells were grown in LB medium supplemented with kanamycin (50 μg/mL), induced with 0.5 mM IPTG at OD_600_ 0.5-0.7, followed by 18h culture at 25 °C prior to harvesting.

### Protein purification

To a resuspension of harvested cells in buffer A (50 mM Tris, 300 mM NaCl, pH 8.0) protease inhibitor cocktail (Pierce™, Thermo Scientific) was added. The mixture was sonicated (15 mins; 5 sec on/5 sec off) and then centrifuged at 17,000 *g* for 40 mins. The soluble protein in supernatant was rocked with a cobalt resin (Thermo Scientific™ HisPur) overnight, then the resin was washed with 5-10 column volumes of buffer A containing 5 mM imidazole. The protein was eluted with 350 mM imidazole in 20 mM Tris, 150 mM NaCl, pH 8.0 buffer. The eluted samples were concentrated by centrifuging at 14,000 *g* for 20 mins using Amicon Ultra Centrifugal Filters (Sigma Aldrich) followed by size exclusion chromatography (ACTA Gold, Cytiva Superdex™ 75 Increase column of 10X300) using a flow rate of 0.8 mL/min. Peak fractions from ∼15 mL were collected, concentrated (HoxA9HD to 1 mg/mL, Pbx1HD to 3 mg/mL), aliquoted, snap-frozen and stored at -80 °C.

### EMSA experiments

To 3 μL of 10X assay buffer (200 mM HEPES, 500 mM NaCl, 50 mM MgCl_2_, 50% glycerol and 2 mM EDTA, pH 8), 3 μL 0.1 mg/mL BSA and 3 μL 0.1% Triton-X, water, protein and the DNA oligonucleotide were added to a total 30 μL reaction volume. The assay mixture was incubated at room temperature for 30 mins followed by addition of 3 μL of 10X native loading dye. 10 μL of this reaction mixture was then loaded into an 8% polyacrylamide gel which was prepared using a recipe (4.4 mL of ddH_2_O, 2 mL of 40% polyacrylamide, 2.4 mL 0.5X Tris-borate-EDTA (pH 8.3), 30 µL 100% Glycerol and 4 µL tetramethylethylenediamine) adapted from earlier.^8,35^ The electrophoresis was conducted on ice at 75 V for 1.5-2 hrs. The gel was washed with water for 15 mins to remove the loading dye, stained with SYBR safe (1:20,000 dilution in ddH_2_O) while gently shaking at room temperature for 30 min, and washed again twice with water for one minute before imaging (ChemiDoc MP, multichannel, blue light excitation). The staining step was skipped for the experiments with the FAM-labelled DNA. During the inhibitor assessment, Pbx1HD was pre-incubated with the inhibitor for 30 mins.

### K_D_ determination with EMSA

The free-DNA signal in each lane was analyzed for area and density using ImageJ.^36^ The fraction of bound DNA (θ_i_) was calculated using the following equation (θi = 1-Ii*/*Io) as described earlier.^35^ The data were analyzed using Microsoft Excel, and curve fitting was performed using the Solver add-in.

### FP assay

The FP experiments were performed in round-bottom, black, 384-well plates (Corning 4514). The assay buffer was 1/10 diluted from 10X buffer (10 mM HEPES, 100 mM NaCl, 5% glycerol, 2 mM EDTA, pH 8.0). The FP signal was measured (Ex/Em: 485/530) using a microplate reader (TECAN Spark). The *K*_D_ was calculated using GraphPad Prism 10.

#### Saturation binding experiments

To a 5 µL solution of serially diluted Pbx1HD, 5 µL of 20 nM FAM-labelled DNA was added, rocked for 1hr at room temperature before measuring the FP.

#### Competitive binding experiment with 6-His peptide

To 5 µL solution of serially diluted peptide in assay buffer, 5 µL of 210 nM Pbx1HD was added. The mixture was incubated at room temperature for 15 mins before adding 5 µL of 30 nM FAM-labeled DNA. The plate was shaken at room temperature for 1h prior to measuring the FP. Final concentrations of 70 nM Pbx1HD, 10 nM FAM-labeled DNA and 100-0.78 µM 6-His peptide were used in a total 15 µL assay-volume.

## ASSOCIATED CONTENT

### Supporting Information

The Supporting Information is available free of charge at https://pubs.acs.org/

The original unedited images of Figure 2 (Figure S1); SYBR-safe stained EMSA replicates of Pbx1HD complex with unlabeled-DNA (Figure S2); EMSA replicates of Pbx1HD complex with FAM-labeled DNA (Figure S3); Saturation binding curve of FAM-labeled DNA with Pbx1HD using FP assay (Figure S4); Ternary DNA complex of Pbx1HD and HoxA9HD with 25 nM unlabeled DNA (Figure S5).

## AUTHOR INFORMATION

### Authors

Kamal Rai – C. Eugene Bennett Department of Chemistry, West Virginia University, Morgantown, West Virginia 26505, United States

Ibukunoluwa Abigail Olaosebikan – C. Eugene Bennett Department of Chemistry, West Virginia University, Morgantown, West Virginia 26505, United States

Pegah Norouzi – C. Eugene Bennett Department of Chemistry, West Virginia University, Morgantown, West Virginia 26505, United States

Gabriella Pettipas – C. Eugene Bennett Department of Chemistry, West Virginia University, Morgantown, West Virginia 26505, United States

Abra G. Dadum – C. Eugene Bennett Department of Chemistry, West Virginia University, Morgantown, West Virginia 26505, United States

Madison Short – C. Eugene Bennett Department of Chemistry, West Virginia University, Morgantown, West Virginia 26505, United States

Kevin C. Courtney – Department of Biochemistry and Molecular Medicine, West Virginia University, Morgantown, West Virginia 26506, United States; Department of Molecular and Cellular Biology, University of Guelph, Guelph, Ontario N1G 2W1, Canada

### Author Contributions

Kamal Rai: Conceptualization, methodology, validation, formal analysis, investigation, writing - original draft; Ibukunoluwa Abigail Olaosebikan: Assisted with HoxA9 protein expression and purification; Pegah Norouzi: Assisted with Pbx1 protein expression; Gabriella Pettipas: Assisted with plasmid transformation; Abra G. Dadum: Assisted with peptide inhibitor synthesis; Madison Short: Assisted with EMSA experiments; Kevin C. Courtney: Review and editing, supervision. Hacer Karatas Bristow: Methodology, supervision, writing - review & editing, project administration. All authors have read, commented on, and agreed to the submitted version of the manuscript.

### Notes

The authors declare no competing financial interest.

## Abbreviations

TF: Transcription factors
EMSA: Electrophoretic Mobility Shift Assay
Pbx1: Pre-B-cell leukemia homeobox-1
HD: homeodomain
LB: Luria-Bertani
FAM: Fluorescein amidite

## Table of Contents Graphic

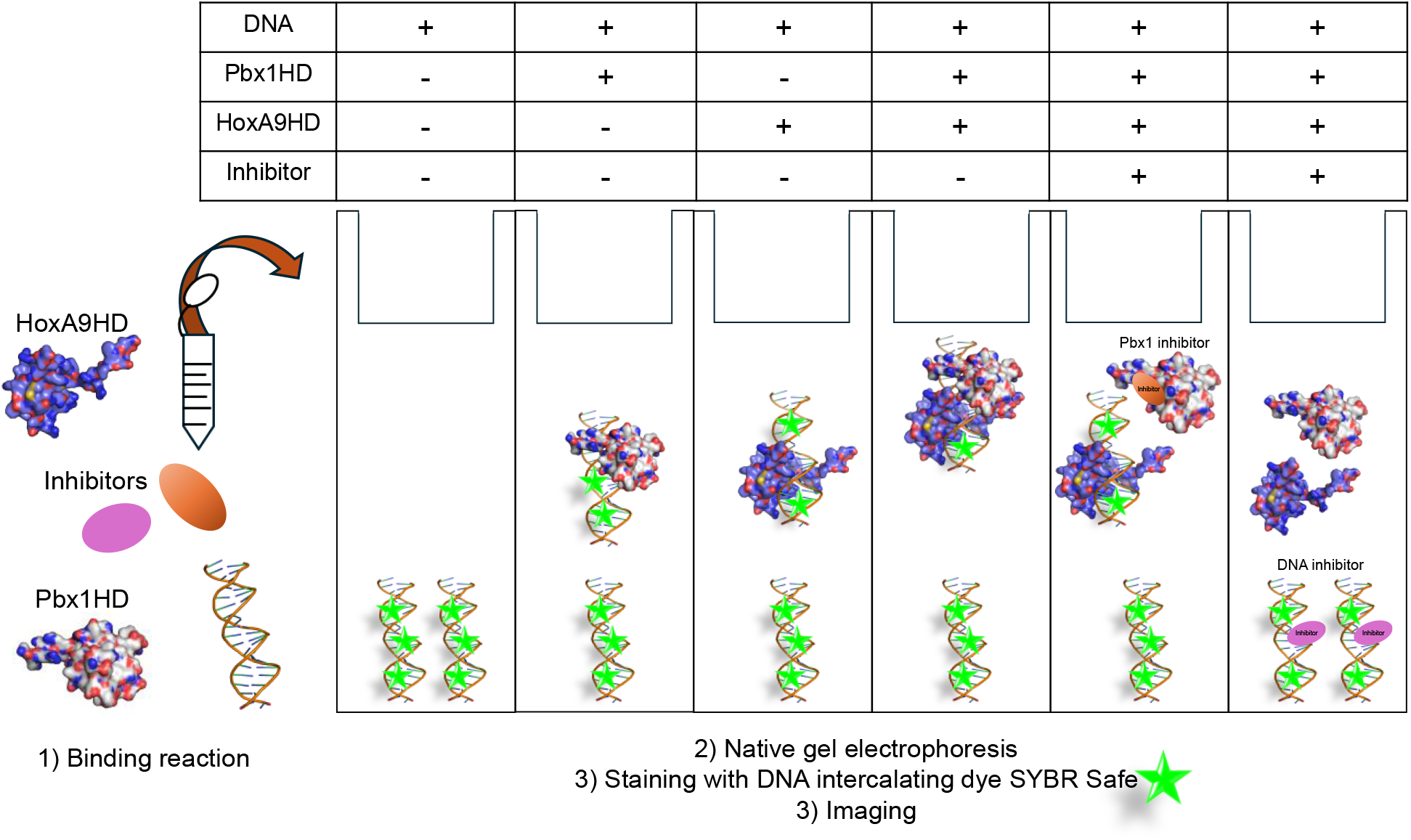

