## Supporting Information for "Development of a Staining-based Electrophoretic Mobility Shift Assay for Analyzing Pbx1, DNA and HoxA9 Interactions"

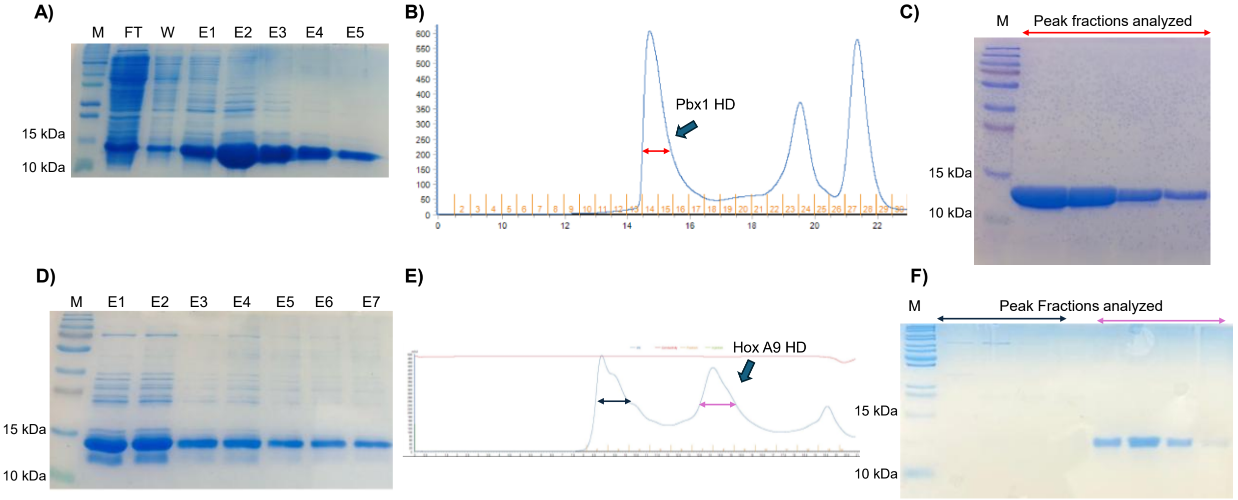


**Figure S1.** The original unedited images of Figure 2. Two-step purification of A-C) Pbx1HD (11.38 kDa) and D-F) HoxA9HD (11.71 kDa). A and D: SDS Page images of affinity purification of Pbx1HD and HoxA9HD, respectively. B and E: Size exclusion chromatograms indicating the collected peak. C and F: SDS Page analysis of peaks collected during SEC. M: Marker, FT: Flow Through, W: Wash, E: repeated elution with 300 mM imidazole.

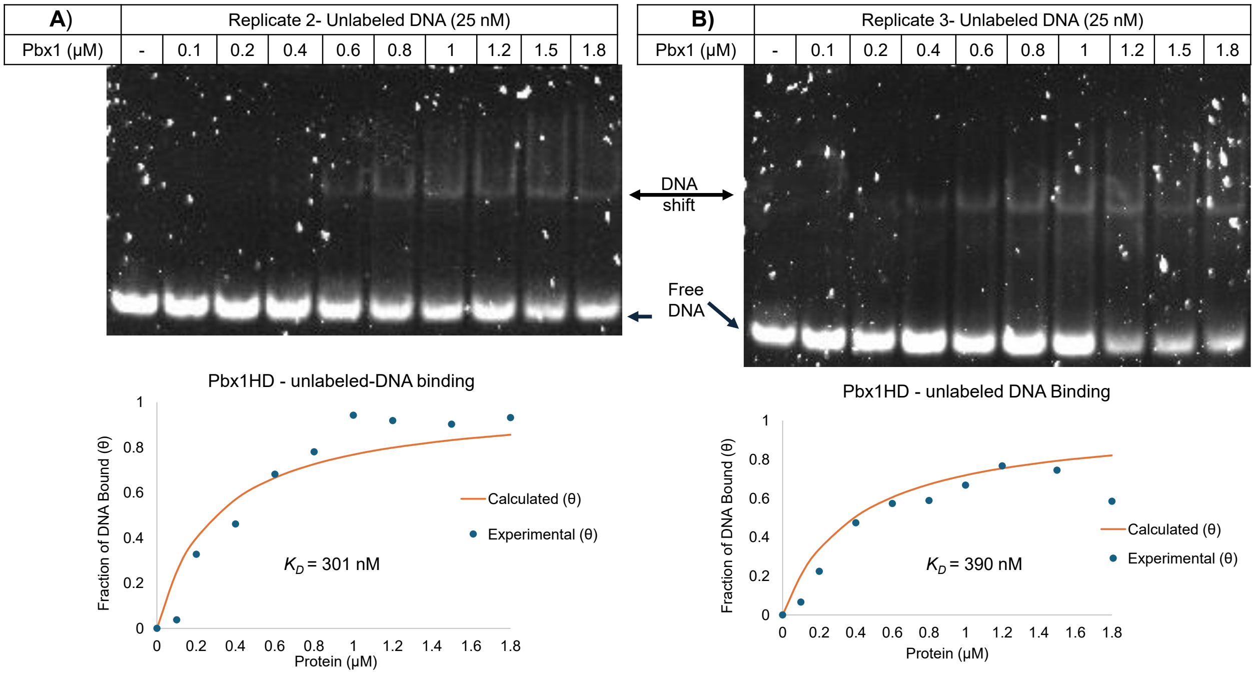


**Figure S2: SYBR-safe stained EMSA replicates of Pbx1HD complex with unlabeled-DNA.** EMSA gel images showing unlabeled DNA stained with SYBR safe on the top and graphs showing the binding curve with their apparent $K_D$ values on the bottom. The free DNA signal on each EMSA gel, as calculated using ImageJ, subtracted from no protein control to determine the fraction of DNA bound (θ) and was plotted against the protein concentration. The apparent $K_D$ values were determined using Langmuir's isotherm.

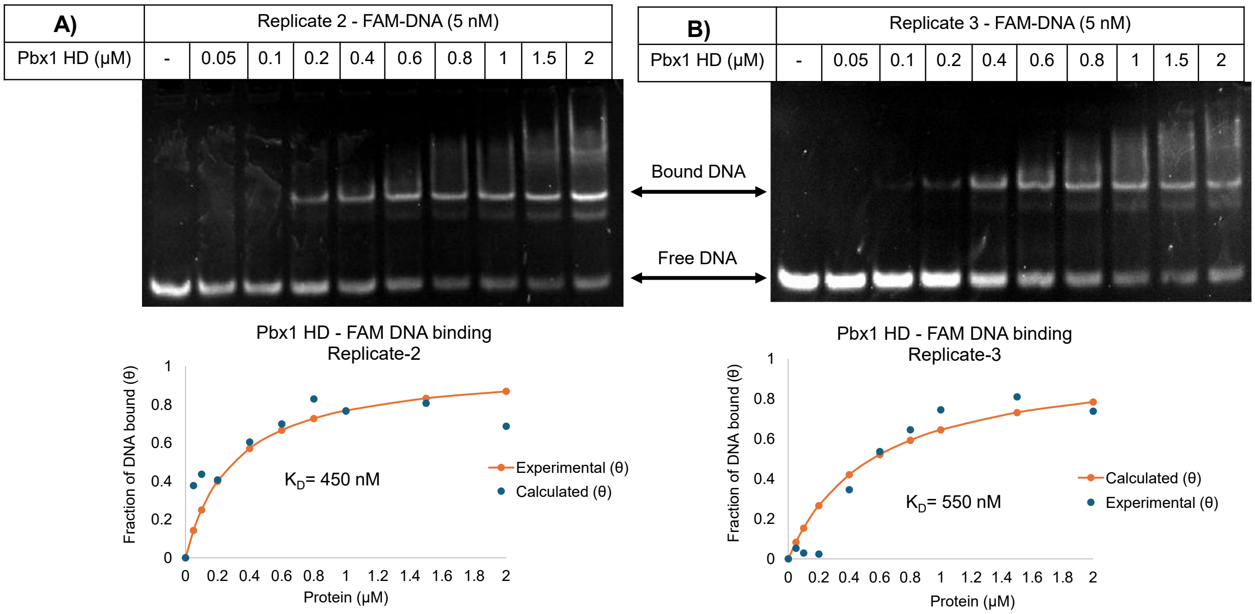


**Figure S3: EMSA replicates of Pbx1HD complex with FAM-labeled DNA.** EMSA gel images showing the FAM-DNA on the top and graphs showing the binding curve with their apparent $K_D$ values on the bottom. The free DNA signal on each EMSA gel, as calculated using ImageJ, subtracted from no protein control to determine the fraction of DNA bound (θ) and was plotted against the protein concentration. The apparent $K_D$ values were determined using Langmuir's isotherm.

| | Biological Replicate 1 | Biological Replicate 2 | Biological Replicate 3 |
|---|---|---|---|
| Best-fit values | | | |
| Bottom | 30.44 | 34.25 | 35.13 |
| Top | 247.8 | 242.7 | 220.4 |
| LogEC50 | -1.128 | -1.150 | -0.9742 |
| HillSlope | 0.7155 | 0.6795 | 0.8031 |
| EC50 | 0.07454 | 0.07080 | 0.1061 |

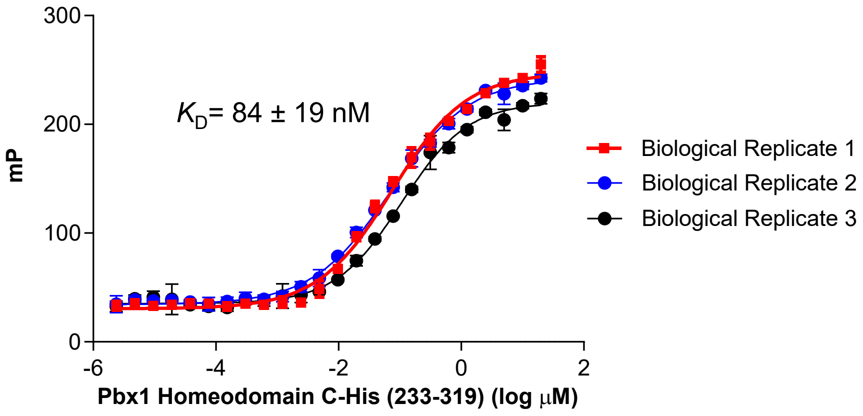


**Figure S4: Saturation binding curve of FAM-labeled DNA with Pbx1HD using FP assay.** FAM-labelled DNA (5 nM) was titrated against Pbx1HD and imaged at 1 hour.

| | | | | |
|---|---|---|---|---|
| DNA (***25 nM***) | + | | | |
| Pbx1HD (0.5μM) | - | + | - | + |
| HoxA9 (0.5μM) | - | - | + | + |

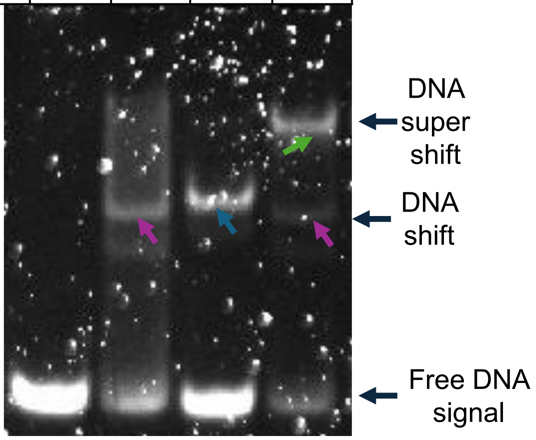


**Figure S5. Ternary DNA complex of Pbx1HD and HoxA9HD with 25 nM unlabeled DNA.** Colored arrows represent the binary complexes of Pbx1HD-DNA (pink), HoxA9HD-DNA (blue) and ternary complex of Pbx1HD-DNA-HoxA9HD (green).
